# Diffusion rather than IFT provides most of the tubulin required for axonemal assembly

**DOI:** 10.1101/268573

**Authors:** J. Aaron Harris, Julie C. Van De Weghe, Tomohiro Kubo, George B. Witman, Karl F. Lechtreck

## Abstract

Tubulin enters the cilia by diffusion and motor-based intraflagellar transport (IFT). The respective contributions of each route in providing tubulin for axonemal assembly are unknown. To attenuate IFT-based transport, we expressed modified GFP-tubulins in strains possessing IFT81 and IFT74 with altered tubulin binding sites. E-hook deficient GFP-β-tubulin normally incorporated into the axonemal microtubules; its transport frequency was reduced by ~90% in control cells and essentially abolished when expressed in a strain possessing IFT81 with an incapacitated tubulin-binding site. Despite the strong reduction in IFT, the share of E-hook deficient GFP-β-tubulin in the axoneme was only moderately reduced indicating that most axonemal tubulin (~80%) enters cilia by diffusion. While not providing the bulk of axonemal tubulin, we propose that motor-based IFT is nevertheless critical for ciliogenesis because it ensures high concentrations of tubulin near the ciliary tip promoting axonemal elongation.

## Introduction

Tubulin is the main protein of the axoneme, the structural core of cilia and flagella (terms used interchangeably; (Borisy and Taylor, 1967)). The axonemal microtubules provide the scaffold onto which dynein arms and other complexes serving in ciliary motility assemble and serve as tracks for intraflagellar transport (IFT). During ciliary assembly, vast amounts of tubulin move from the cell body into the organelle, e.g., a 12-μm long axoneme contains ~350,000 tubulin dimers (Kubo et al., 2016). Tubulin enters cilia by diffusion and intraflagellar transport (IFT; (Craft et al., 2015; Hao et al., 2011; Luo et al., 2017)). In IFT, arrays of IFT particle (= IFT trains) travel bidirectionally along the axonemal microtubules by the means of motor proteins (Cole et al., 1998; Kozminski et al., 1993). While IFT speed and frequency remain essentially unaltered, the frequency of tubulin transport is upregulated while cilia elongate and attenuated in non-growing cilia. Similar to other small soluble proteins, tubulin continuously diffuses into cilia apparently moving freely through the ciliary gate formed by the transition zone at the ciliary base (Craft et al., 2015; Harris et al., 2016; Kee and Verhey, 2013). The respective contributions of diffusion and IFT in supplying tubulin for axonemal assembly are unknown. Manipulating IFT-tubulin interactions could help to estimate the share of tubulin supplied by each path.

*In vitro* interaction studies revealed that the N-terminal domains of the IFT particle proteins IFT81 and IFT74 form a bipartite tubulin-binding module (Bhogaraju et al., 2013). In detail, the calponin homology (CH)-domain of IFT81 binds to the globular core of the αβ-dimer providing specificity and the basic N-terminus of IFT74 interacts with the acidic C-terminal E-hook of β-tubulin stabilizing the interaction. *In vivo*, the deletion of the N-terminal 130 residues of IFT74 (IFT74Δ130) severely decreased the speed of ciliary regrowth; ciliary length, however, was only moderately reduced (Brown et al., 2015). Mutations of two or five basic residues in the CH-domain of IFT81 diminishes tubulin binding *in vitro* (Bhogaraju et al., 2013). However, *C. reinhardtii* expressing an altered IFT81 with five such point mutations (IFT81-5E) still assembles near full-length flagella albeit at a reduced rate (Kubo et al., 2016). In both the IFT81-5E and IFT74Δ130 strains IFT transport of GFP-α-tubulin was reduced potentially explaining the delay in ciliary growth. Double mutants in the tubulin binding sites of IFT81 and IFT74 assemble none or severely truncated flagella with an otherwise normal ultrastructure supporting the notion that the IFT81/74 module promotes ciliogenesis by facilitating IFT transport of tubulin (Kubo et al., 2016). While great care was taken to ensure that the above-described manipulations of IFT81 and IFT74 are specifically affecting tubulin transport, defects in IFT itself cannot be excluded. The strain expressing the N-terminally truncated IFT74, for example, displayed anomalies in IFT speed and frequency and the abundance of various IFT proteins in the cell body; further, phospholipase D was enriched in cilia indicative for defects in retrograde IFT (Brown et al., 2015; Lechtreck et al., 2013). Due to the short length of the cilia, IFT could not be assessed in the double mutant. It is therefore of interest to analyze how changes in tubulin itself affect its transport by IFT. Here, we focused on the E-hook of β-tubulin, which has been predicted to interact with the N-terminal domain of IFT74.

In the ciliate *Tetrahymena thermophila*, genomic replacement of wild-type β-tubulin with versions lacking a functional E-hook severely impairs cell viability and ciliary assembly (Duan and Gorovsky, 2002; Xia et al., 2000). Thus, approaches altering the entire tubulin pool of a cell are unsuited to determine the specific effect of such manipulations on tubulin transport by IFT. Here, we expressed GFP-tagged α- and β-tubulins in *C. reinhardtii* to levels of 5-10% of the total tubulin and analyzed how the altered E-hooks affect IFT transport without perturbing either IFT itself, IFT of the endogenous tubulin, or ciliary assembly. This approach has the added potential to shed light on the relative contributions of IFT and diffusion in ciliary tubulin supply by analyzing how a reduction in IFT of a specific assembly-competent GFP-tagged tubulin will affect its share in the axoneme.

We show that the deletion of the E-hook of β-tubulin severely reduced its ability to bind to IFT trains; the effect is specific for the E-hook of β-tubulin. IFT transport of E-hook-deficient β-tubulin was nearly abolished in a strain expressing carrying IFT81 with defects in its tubulin-binding CH-domain. Nevertheless, E-hook deficient β-tubulin remained abundant in the axoneme suggesting a smaller than anticipated role of IFT in providing tubulin for axonemal assembly. Based on our observations, we estimate that about 20% of the total tubulin in the axoneme is provided by IFT while ~80% enter cilia by diffusion. We postulate that the elongated geometry of cilia with just a narrow opening to the cell body delays tubulin entry by diffusion requiring tubulin transport by IFT to ensure ciliary assembly.

## Results

### The E-hook of β-tubulin promotes tubulin-IFT interaction in vivo

SuperfolderGFP (GFP) was fused to the N-terminus of β-tubulin and expressed in wild-type *Chlamydomonas reinhardtii* (Fig. 1). Western blotting using anti-GFP and anti-β-tubulin confirmed expression and the presence of GFP-β-tubulin in flagella (Fig. 1A). Life cell imaging showed that GFP-β-tubulin is incorporated into both cytoplasmic and axonemal microtubules (Fig. 1B). As previously reported for GFP-α-tubulin, IFT of GFP-β-tubulin was rarely observed in steady-state flagella but occurred with high frequency in elongating flagella obtained by first deciliating the cells via a pH shock. (Fig. 1C) (Craft et al., 2015). Therefore, the transport frequencies for all tubulin constructs used in this study were obtained using regenerating flagella with a length of ~4 – 9 μm. A comparison of strains expressing different amounts of tagged β-tubulin revealed an approximately linear correlation between the total amount of GFP-β-tubulin expressed, the frequency of its transport in regenerating flagella by IFT, and its share in the axoneme (Fig. 1D, E).

**Figure 1).**
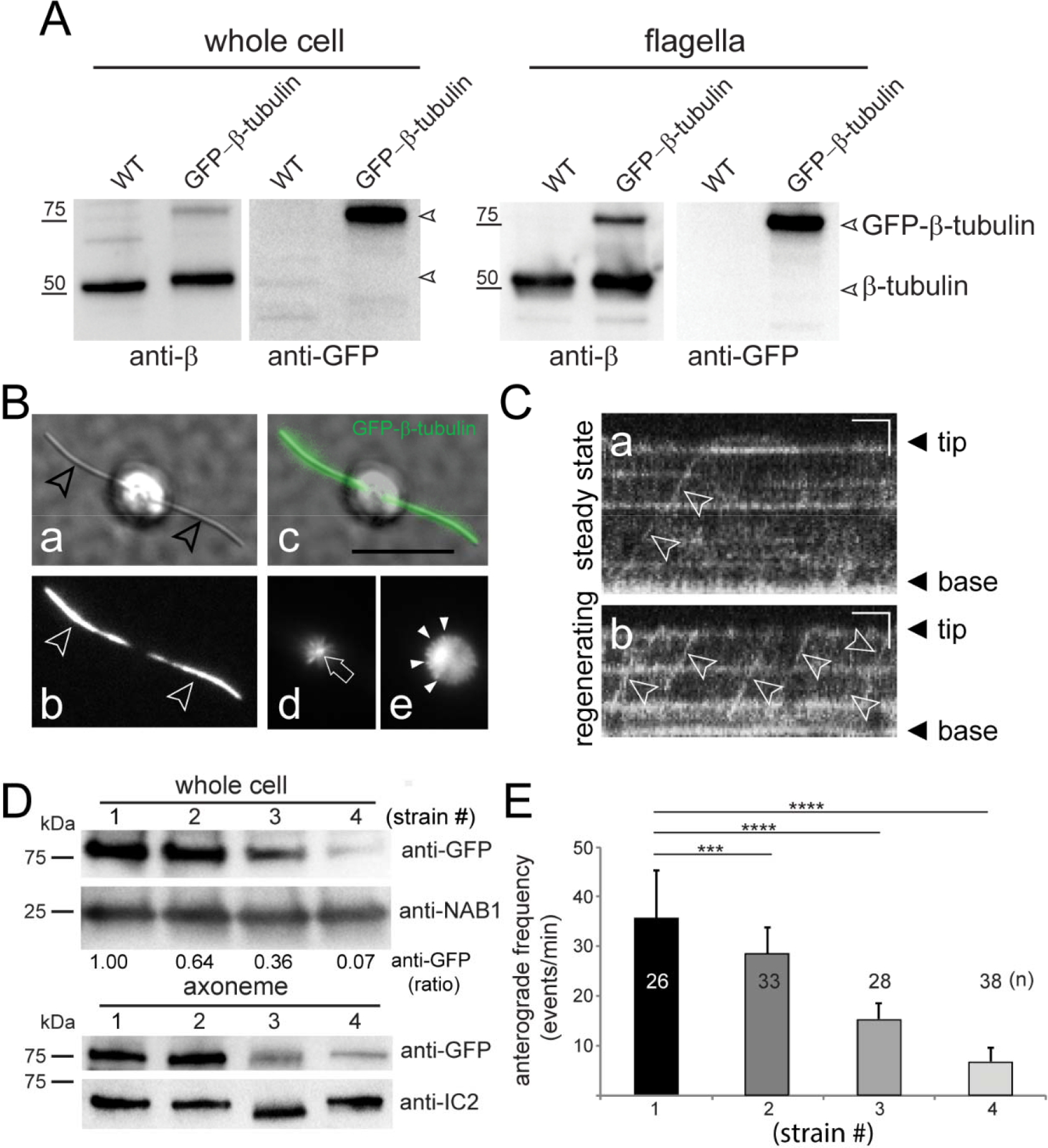
GFP-β-tubulin is transported by IFT and incorporated into microtubules. A) Western blots analyzing whole cells and isolated flagella from a wild-type strain expressing GFP-β-tubulin and an untransformed control strain (WT). Western blots were stained with apolyclonal antibody to β-tubulin and anti-GFP; the position of endogenous β-tubulin, the GFP tagged version, and marker proteins are indicated. B) Life cell imaging of a cell expressing GFP-β-tubulin in DIC (a) and TIRF (b, d, e); an overlay image is shown in c. The TIRF images present a focal series showing the level of the cilia (marked by arrowheads in b), the basal bodies (arrow in d), and a more posterior section showing the cortical microtubules of the cell body (arrowheads in e). Bar = 10 μm. C) Kymograms showing IFT transport (arrowheads) of GFP-β-tubulin in steady-state (a) and regenerating (b) flagella. The tip and the base of the cilia are indicated. Bars = 2 s and 2 μm. D) Western blot of whole cell and axoneme samples obtained from four wild-type strains (Nos. 1 – 4) expressing different levels of GFP-β-tubulin. Blots were stained with anti-GFP; antibodies to the cell body protein NGFI-A binding protein 1 (NAB1) and the axonemal dynein intermediate chain 2 (IC2) were used as loading control for the cell body and axonemes, respectively. E) Frequency of GFP-β-tubulin anterograde transport during flagellar regeneration in the strains shown in Fig. 1D.

To investigate the role of the βE-hook for IFT of tubulin, GFP-tagged β-tubulin was modified by deleting of the C-terminal 13 residues (ΔE-hook), replacing five glutamic acid residues in the E-hook with alanine undermining charged-based interactions (−5E-hook), or substituting the E-hook of β-tubulin with that of α-tubulin (αE-hook; Fig. 2, Table S1). Cells expressing the transgenic GFP-tubulins grew normally, showed wild-type motility, assembled full-length flagella at normal rates, and IFT of tubulin progressed at standard velocities (Table S1, not shown). To determine whether the E-hook modifications affected the frequency of GFP-β-tubulin transport, we selected strains expressing similar amounts of the transgenes as determined by Western blotting of whole cell samples (Fig. 2A). For all four constructs, ~8% of the total GFP-tagged protein was in the flagella and ~90% of the flagellar GFP-tubulin was in the axonemal fraction indicating that the fusion proteins entered the flagella and were assembly competent (Fig. 2B). All three mutations in the βE-hook resulted in significant reductions in the frequency of anterograde tubulin IFT with the E-hook deficient construct being the most affected (2.4 events/min, SD 2.9 events/minute, n=98 flagella analyzed compared to 21 events/minute, SD 14.3 events/minute, n=85 for wild-type β-tubulin; Fig. 2C).

**Figure 2).**
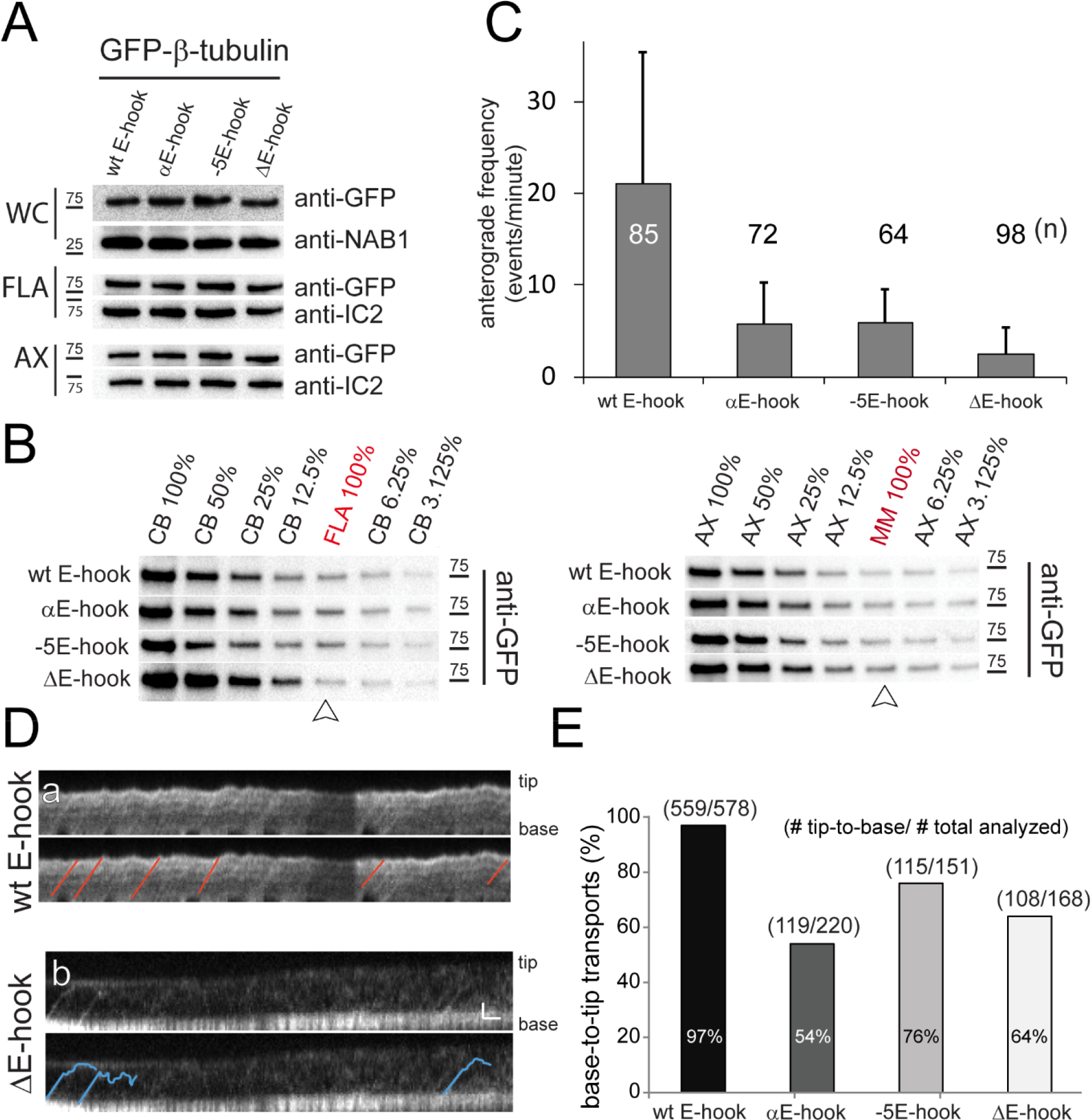
The E-hook of β-tubulin promotes transport by IFT. A) Western blot analyses of whole cells (WC), isolated flagella (FLA), and axonemes (AX) from strains expressing GFP-β-tubulin with wild-type and modified E-hooks. Anti-GFP was used to visualize the tagged β-tubulins and antibodies to NAB1 and IC2 as loading controls for WC and FLA/AX, respectively. B) Left side: Western blots comparing the amount of GFP-β-tubulin in the deflagellated cell bodies (CB) to that in flagella (FLA; left) and (right side) comparing its level in the axonemes (AX) to that in the detergent soluble membrane+matrix fraction (MM, right) using a dilution series. 100% indicates that equivalents of the two fractions were loaded (i.e., one CB per two FLA etc.). C) Histogram showing the average frequency of anterograde IFT events observed for the GFP-tagged β-tubulins. The number of regenerating cilia analyzed for each strain (n) and the standard deviation are indicated. D) Kymograms showing transport of full-length and E-hook deficient (ΔE-hook) β-tubulin in regenerating flagella. Selected tracks are marked on the duplicates (bottom panels, respectively). Note the reduced frequency and reduced run length of transports involving the truncated GFP-β-tubulin (blue lines). Bar = 1s 1μm. E) Histogram comparing the share (in %) of early-terminating transport events of wild-type and modified GFP-β-tubulins. The number of processive base-to-tip events and the total number of events analyzed are indicated.

Loss of the βE-hook weakens the interaction of tubulin with IFT in vitro (Bhogaraju et al., 2013) and we wondered how loss of the βE-hook affects the processivity of tubulin transport in cilia. The axonemal protein DRC4-GFP, for example, frequently (~40%) dissociates from IFT trains before arriving at the flagellar tip (Wren et al., 2013). In contrast, IFT transport of GFP-α-tubulin mostly (~98%) proceeds in one run to the tip indicative for a stable interaction with IFT trains (Craft et al., 2015). Similarly, 97% (n= 559) of the transports involving wild-type GFP-β-tubulin proceeded nonstop to the tip of the elongating flagella whereas such processive IFT transports were reduced by 25-45% for the constructs with modified E-hooks (Fig. 2D and E). In those strains, GFP-β-tubulin was observed converting from IFT transport to diffusion indicative for a dissociation from IFT (Fig. 2D).

Considering the apparent importance of the βE-hook for tubulin binding to IFT, we transplanted the E-hook and neighboring regions of β-tubulin onto GFP or mNeonGreen (mNG; Fig. S1). Similar to GFP alone, the GFP/mNG-βΔE-hook fusion proteins readily entered flagella by diffusion. In regenerating flagella, IFT transport of GFP alone was not observed. Anterograde IFT was observed for the GFP/mNG-βE-hook fusions albeit at extremely low frequencies (<0.05 events/min) indicating an almost complete absence of binding to IFT trains (Fig. S1). In summary, the βE-hook is neither sufficient nor necessary for IFT transport; its loss, however, reduces the frequency and processivity of tubulin transport supporting the view that the βE-hook stabilizes IFT-tubulin interactions.

### High frequency transport of tubulin requires the E-hook of beta-tubulin to interact with IFT74

Bhogaraju et al. (2013) proposed that tubulin binding by IFT trains involves the CH-domain of IFT81 and an interaction between the E-hook of β-tubulin and the N-terminal domain of IFT74 (Fig. 3A). To further investigate the validity of this model, we analyzed tubulin transport in *ift81-1 IFT81-5E*, a strain in which five basic residues (K73, R75, R85, K112 and R113) critical for tubulin binding by the IFT81 CH-domain were replaced by glutamates, and in *ift74-2 IFT74ΔN1-130*, an *IFT74* null mutant rescued with a version of IFT74 lacking the proposed N-terminal tubulin-binding domain (Fig. 3B-G) (Brown et al., 2015; Kubo et al., 2016).

**Figure 3).**
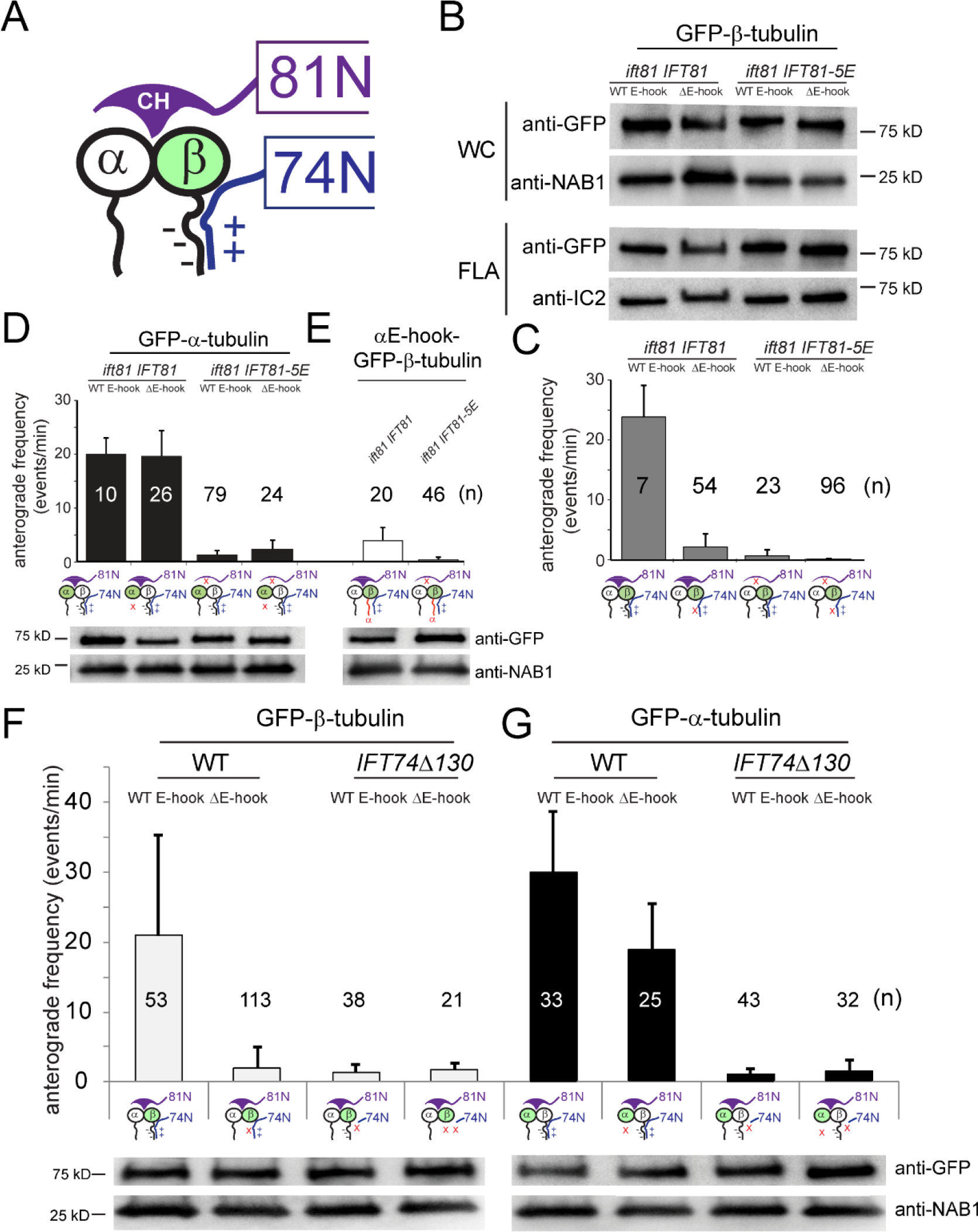
Transport of E-hook deficient β-tubulin is sensitive to mutations in IFT81. A) Schematic presentation of tubulin binding by the IFT81N/IFT74N module (modified after (Bhogaraju et al., 2013)). B) Western blot comparing the amounts of wild-type and ΔE-hook GFP-β-tubulin in the *ift81-1 IFT81* and the *ift81-1 IFT81-5E* rescue strains. The transgene expression levels were similar in all four strains. For this blot, loading was adjusted to result in anti-GFP signals of similar strength. C) Histogram showing the average frequencies of anterograde IFT events observed for full-length (wt) and truncated (Δβ) GFP-tagged β-tubulins in cilia of the *ift81-1 IFT81* and the *ift81-1 IFT81-5E* strains. The number of regenerating cilia analyzed for each strain (n) and the observed standard deviation are indicated. D) Frequency of anterograde IFT for full-length and E-hook deficient GFP-α-tubulin in the *ift81-1IFT81* and the *ift81-1 IFT81-5E* strains. The corresponding western blots of the whole cell samples of these strains are shown below the graph. E) Anterograde transport of αE-hook-GFP-β-tubulin (see D for details). F, G) Anterograde transport of full-length and E-hook deficient GFP-β-tubulins (F) and GFP-α-tubulin (G) in the *ift74-2 IFT74Δ130* strain and wild-type controls. Western blots of whole cell documenting similar expression of the transgenes are shown (bottom).

E-hook deficient β-tubulin is predicted to be unable to interact with the N-terminal domain of IFT74 and thus its binding to IFT trains should depend on the CH-domain of IFT81. In the *ift81-1 IFT81-5E* background, intermediate transport frequencies were observed for full-length β-tubulin while anterograde IFT of ΔE-hook β-tubulin was essentially abolished (<99% compared to full-length β-tubulin in strains with wild-type IFT81; Fig. 3B, C). Sporadic transport of E-hook deficient β-tubulin in *ift81-1 IFT81-5E* could be due to residual binding by IFT81-5E or low capacity binding elsewhere on IFT. The *ift74-2 IFT74ΔN1-130* strain lacks the proposed βE-hook-binding site of IFT74. The anterograde IFT frequencies of full-length and E-hook deficient β-tubulin were similarly low suggesting that both bind with similar strength to IFT trains presumably via the remaining IFT81-CH site (Fig. 3F). Thus, the lack of the N-terminal region of IFT74 renders IFT unable to discriminate between full-length and E-hook deficient β-tubulin.

We wondered whether IFT transport specifically requires the E-hook of β-tubulin or whether the E-hook of α-tubulin also contributes to IFT. In wild-type IFT cells, transport of E-hook deficient α-tubulin was somewhat reduced but still occurred at a high frequency when compared to E-hook deficient β-tubulin (Fig. 3D, G). Also, strains expressing *ift74-2 IFT74ΔN1-130* or *ift81-1 IFT81-5E* transported full-length and E-hook deficient α-tubulin with similar intermediate frequencies (Fig. 3D, G). Thus, the IFT system is largely insensitive to the loss of the αE-hook. Further, substitution of the E-hook of GFP-β-tubulin with that of α-tubulin did not restore normal transport frequencies in wild-type (Fig. 2C) or the *ift81-1 IFT81-5E* background (Fig. 3E). The data support the model by Bhogaraju et al. (2013) in which the E-hook of β-tubulin interacts with the N-terminal domain of IFT74. We further show that loss of βE-hook in combination with mutations in the IFT81-CH domain severely reduce IFT transport of GFP-tubulin.

### Near abrogation of IFT of tagged β-tubulin only moderately reduces its axonemal presence

In addition to IFT, tubulin enters flagella also by diffusion. The expression of a tagged assembly-competent tubulin with strongly decreased binding to IFT facilitates an assessment of the respective contributions of IFT and diffusion in supplying axonemal tubulin. If, for example, ~90% of the flagellar tubulin were delivered by IFT, one would expect that a strong reduction in IFT of a specific tubulin species will be reflected by an almost equally strong reduction of its share in the axoneme. Toward this end, we analyzed the correlation between the frequency of IFT and the axonemal share for full-length and E-hook deficient GFP-β-tubulin (Fig. 4). Loss of the βE-hook reduced the frequency of IFT by ~90% in wild-type, the *ift81* IFT81 rescue strain (expressing a wild-type IFT81 transgene), and the *ift81* IFT81-5E rescue strain, each in comparison to full-length GFP-β-tubulin (Figs. 4A). Western blot analyses of axonemes, however, documented only a moderate reduction of E-hook deficient GFP-β-tubulin compared to the full-length version (Figs. 2A, 4B). Quantitative analysis of western blots showed a reduction of in average ~10% (SD 33%, n = 9) for truncated vs. full-length GFP-β-tubulin from nine independent flagella isolates (four, three, and two in the wild-type, *ift81* IFT81, and *ift81* IFT81-5E backgrounds, respectively; Figs. 4C, S2A). Western blots of isolated axonemes using anti-β-tubulin confirmed that full-length and truncated GFP-β-tubulin were present in similar ratios to the endogenous β-tubulin; for this experiment we used a polyclonal anti-*Chlamydomonas*-β-tubulin since most commercial antibodies to β-tubulin react with the E-hook (Fig. 4D; (Silflow and Rosenbaum, 1981)). Even in the near absence of transport by IFT, as observed for E-hook deficient GFP-β-tubulin in the *ift81 IFT81-5E* strain (~99.8% reduction in frequency in comparison to GFP-β-tubulin in wild-type), the truncated GFP-tagged tubulin was still well presented in the flagella (Fig. 4E; ~50% of the wild-type control, n=2 independent isolates). The average reduction of E-hook deficient GFP-β-tubulin’s share in the axoneme was 17% (SD 35%; n=10 flagellar isolates). In live cell imaging, the flagella of strains expressing full-length GFP-β-tubulin (in the *ift20* IFT20-mCherry background to facilitate identification) and E-hook deficient GFP-β-tubulin were of similar brightness (Fig. 4F). In summary, the strong reduction in the frequency of IFT transport observed of E-hook deficient β-tubulin is not matched by a proportional reduction of its presence in the axoneme.

**Figure 4).**
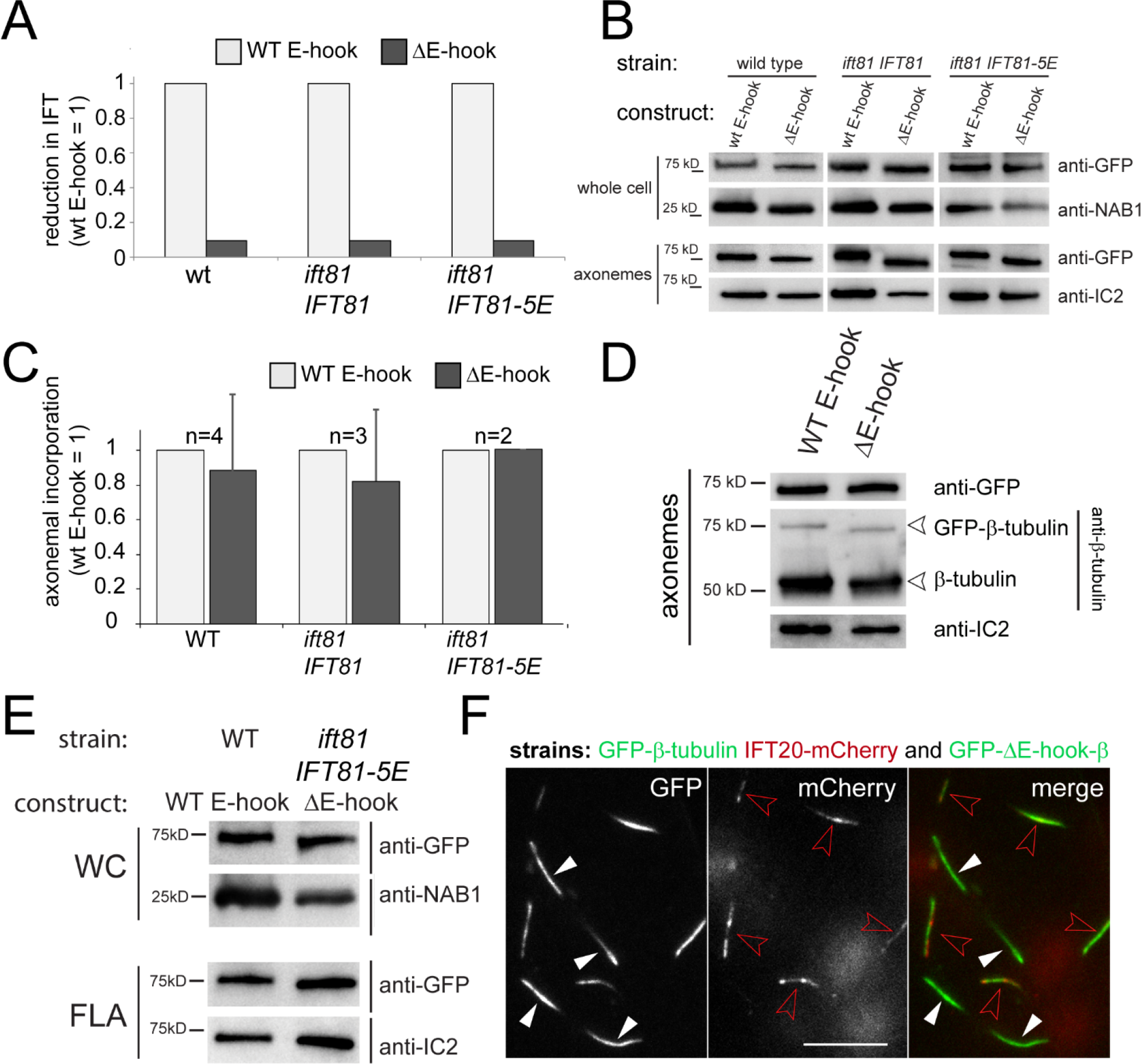
Abated IFT of E-hook deficient tubulin does not cause a proportional reduction of its presence in the axoneme. A) Histograms comparing the reduction in IFT transport of ΔE-deficient vs. full-length β-tubulin in wild-type, *ift81-1 IFT81*, and *ift81-1 IFT81-5E* rescue strains. The frequencies observed for the full-length GFP-β-tubulin in each background were set to 1. See Fig. 3 for the original data, SDs, and n values. B) Western blot of wild-type and *ift81-1* rescue strains comparing whole cell and axoneme samples using anti-GFP and antibodies to NAB and IC2, as loading controls. C) Histograms comparing the changes in the axonemal amounts of ΔE-deficient vs. full-length β-tubulin in wild type, and the *ift81-1 IFT81* and *ift81-1 IFT81-5E* rescue strains as deduced from anti-GFP stained Western blots. The band intensities of the full-length GFP-β-tubulin in each back ground were set to 1. n, number of independent flagella isolates. D) Western blot comparing isolated axonemes from a wild-type strain expressing full-length or truncated β-tubulin stained with anti-GFP, anti-β-tubulin, and anti-IC2. E) Western blot of whole cell (WC) and flagella (FLA) samples comparing strains expressing full-length β-tubulin in wild type vs. ΔE-deficient β-tubulin in the *ift81-1 IFT81-5E* strain side by side. The antibodies used are indicated. F) TIRF images showing the GFP and mCherry signals in cilia of live cells. Cells expressing E-hook deficient β-tubulin and cells expressing full-length β-tubulin in the *ift20-1* IFT20-mCherry background were mixed to allow for a comparison of the signal strength with the same microscope settings. Bar = 10 μm.

Our data suggest a limited quantitative contribution of IFT in providing tubulin for axonemal assembly. However, cells with defects in the IFT81/74 tubulin-binding module largely fail to assemble flagella. We therefore searched for possible alternative mechanisms potentially explaining the surprisingly low effect the reduction in the transport frequency of truncated β-tubulin has on its axonemal share. Mutations in IFT81N or IFT74N reduce IFT of tubulin and slow down flagellar growth suggesting that reduced IFT of tubulin could lower the rate of flagellar elongation (Kubo et al., 2016). Here, we used regenerating flagella to determine the IFT frequency while flagella assembled after the previous cell division were used for our biochemical analysis. The former requires approximately 90 minutes to regenerate flagella of 12-μm length whereas post-mitotic assembly is considerably slower completing ~ 8-μm long flagella in ~4 hours ((Madsen et al., 1983; Rosenbaum et al., 1969); REPLACE MADSEN WITH: Madey and Melkonian, 1990). Postmitotic flagellar assembly could involve less IFT and more diffusion of precursors, potentially explaining the high share of ΔE-hook GFP-β-tubulin in flagella despite its low frequency of IFT. Western blot analysis, however, did not reveal a significant difference in the share of full-length and E-hook deficient β-tubulin in postmitotic vs. regenerated axonemes (Fig. S2B). Further, a strong discrimination of truncated GFP-β-tubulin by retrograde IFT could enrich it relative to the full-length protein. The frequencies of retrograde tubulin traffic, however, were negligibly low compared to anterograde traffic and were similar in range for full-length GFP-β-tubulin and its derivates (Fig. S2C, D). Also, the total amount of GFP-β-tubulin tolerated in axonemal microtubules could be limited and even low IFT frequencies as observed for truncated GFP-β-tubulin could still be sufficient to saturate GFP-tubulin incorporation. We consider this situation unlikely because the IFT frequency and axonemal quantity of full-length GFP-β-tubulin correlated well over a wide range of expression levels (Fig. 1D, E). Finally, E-hook deficient β-tubulin could incorporate into the axoneme at a significantly higher rate than the full-length protein. Of note, subtilisin-treated E-hook deficient tubulin has an about 50X lower critical concentration for polymerization into rings and sheets albeit assembly of bona fide microtubules was not observed (Bhattacharyya et al., 1985; Sackett et al., 1985). If E-hook deficient β-tubulin indeed incorporates into axonemal microtubule at a much higher rate than the full-length protein, one would expect the truncated version to be more abundant in the axoneme than full-length GFP-β-tubulin under conditions when both proteins show the same frequency of IFT. This is the case in the *ift74-2 IFT74ΔN1-130* background when tubulin transport relies solely on the IFT81-CH. Comparison of the axonemes revealed similar shares for full-length and truncated GFP-β-tubulin expressed at similar levels in *ift74-2 IFT74ΔN1-130* (Fig. S2E). In conclusion, our observations are explained best by assuming that IFT transports only a fraction of the tubulin present in the axoneme and that most tubulin enters flagella by diffusion.

## Discussion

Tubulin enters cilia by diffusion and as a cargo of IFT. The latter is regulated in a ciliary-length dependent manner and this regulation is likely to contribute to ciliary length control (Craft et al., 2015; Hao et al., 2011; Kubo et al., 2016). To explore the respective contributions of each route in providing tubulin for axonemal assembly, we manipulated tubulin transport by IFT taking advantage of the detailed knowledge of IFT-tubulin interactions. Previously *in vitro* studies showed that the E-hook of β-tubulin confers stable binding of tubulin dimers to the IFT81N/74N module by interacting with IFT74 (Bhogaraju et al., 2013). In agreement with this model, removal of the E-hook of GFP-tagged β-tubulin reduced IFT by ~90%. However, the strong reduction in IFT was not mirrored by a comparative reduction of the share of E-hook deficient β-tubulin incorporated into the axoneme. We propose that IFT’s predominate role is to regulate intraciliary tubulin concentrations rather than providing the bulk of tubulin needed for ciliary assembly.

### IFT74 specifically interacts with the βE-hook to promote tubulin IFT

Bhogaraju and colleagues (2013) showed that tubulin-binding by the IFT81N/IFT74N module was significantly reduced when the E-hook of β-tubulin was removed by a brief subtilisin treatment. Subtilisin treatment only mildly affected tubulin binding by IFT81N alone suggesting that the positively charged IFT74N interacts with negatively charged E-hook of β-tubulin. Our *in vivo* observations are in agreement with this model and provide the additional insights: 1) The E-hook of β-tubulin is not essential for binding to IFT trains or incorporation into the axonemal microtubules; it is also not sufficient for IFT transport. 2) IFT74N’s interaction with tubulin is specific for the β-tail while the negatively charged E-hook of α-tubulin is expendable. 3) One of the two predicted tubulin-IFT contacts (i.e., IFT81-CH and tubulin dimers, IFT74N and βE-hooks) is sufficient to maintain intermediate IFT transport frequencies and, as recently shown by Kubo et al. (2016), ciliary assembly. 4) GFP-tubulin transport was essentially abolished when the deletion of the βE-hook was combined with mutations in IFT81N, an arrangement thought to torpedo both interactions of the IFT81N/IFT74N module with tubulin. This suggests that IFT81N/IFT74N module is the major tubulin binding site on anterograde IFT trains and argues against the presence of additional tubulin binding sites independent of the IFT81N/IFT74N module (Bhogaraju et al., 2014). This statement is supported by the recent finding from Kubo et al. (2016) that double mutants in IFT81N and IFT74N are essentially unable to assemble flagella.

### The beta-E-hook in tubulin transport and microtubule stability

Tubulin E-hooks are subjected to a variety of posttranslational modifications, which are thought to be critical for the stability of axonemal microtubules. After genomic replacement of the sole conventional β-tubulin gene of *Tetrahymena thermophila* with versions carrying modified E-hooks, cells assemble shorter than normal cilia frequently lacking the ciliary central pair and the B-tubules of the doublets (Brown et al., 1999; Thazhath et al., 2004; Thazhath et al., 2002; Xia et al., 2000). The mutant phenotype has been explained by the lack of posttranslational modifications, in particular polyglycylation, which occurs on five sites in the *T. thermophila* βE-hook. Both the decrease in tubulin IFT in *C. reinhardtii* and the structural defects in the cilia of *T. thermophila* are specific for changes to the βE-hook raising the question whether reduced IFT of tubulin could contribute to the axonemal defects observed in the *T. thermophila* mutants. Removal of the E-hook from the sperm-specific β2-tubulin inhibits flagellar development in *Drosophila melanogaster*, and its replacement with the β1 E-hook obstructs central pair assembly (Nielsen et al., 2001). *Drosophila* sperm flagella, however, assemble independently of IFT in the cytoplasm; diminished tubulin transport is therefore unlikely to contribute to the fly phenotype. We conclude that the E-hook of β-tubulin is critical for both high frequency IFT of tubulin and the stability of axonemal microtubules.

### The bulk of axonemal tubulin is supplied by diffusion

Here, we circumvented problems in axonemal assembly caused by mutations in βE-hook by co-expressing GFP-tagged tubulin in cells possessing wild-type endogenous tubulins. This approach allowed us to analyze IFT transport of modified tubulins independently of ciliary assembly. Surprisingly, a strong reduction in IFT of E-hook deficient β-tubulin was not matched by a proportional reduction of its share in the axoneme. After assessing several possible mechanisms, we conclude that most of the tubulin in the axoneme enters the cilium by diffusion.

Estimates of the transport capacity of IFT support this notion: While several other IFT proteins possess CH-domains, only the ones of IFT81 and IFT54 bind tubulin (Bhogaraju et al., 2014; Taschner et al., 2016). Recently, it has been shown that the CH-domain of IFT54 is expendable for ciliary assembly (Zhu et al., 2017). Thus, it is likely that the IFT81N/IFT74N module forms or is part of the only high capacity tubulin binding side of IFT. Ultrastructural data indicate that each IFT train contains about 40 IFT-B complexes and approximately 60 IFT trains enter the cilium each minute. The oligomeric state of tubulin during transport is unknown but the low stability of tubulin protofilaments suggests that tubulin is transported as a dimer. Under these assumptions, IFT could transport approximately 25% of the ~10,000 tubulin dimers required each minute during phases of rapid ciliary growth (~350 nm/minutes; (Bhogaraju et al., 2014)). Apparently, the IFT pathway simply lacks the capacity to be the major provider of axonemal tubulin during ciliary assembly. On the other hand, FRAP analysis revealed a continuous high-frequency influx of GFP-tubulin into cilia by diffusion (Craft et al., 2015). The experimental data presented here suggest that roughly 4/5^th^ of the axonemal tubulin enter the cilium by diffusion.

### Ciliary geometry delays diffusion explaining the need for tubulin transport by IFT

The above notion needs to be reconciled with observations indicating a critical role of IFT in providing tubulin for ciliary assembly, e.g., defects in the IFT81N/IFT74N module reduce the ciliary growth rate or impair ciliogenesis (Bhogaraju et al., 2013; Kubo et al., 2016). Apicomplexan parasites such a *Plasmodium* rapidly assemble numerous axonemes within the cell body without IFT (Billker et al., 2002; van Dam et al., 2013). In that situation, diffusion is sufficient to provide the tubulin necessary for axonemal assembly as it can be generally assumed for other large-scale microtubule-based assemblies such as the mitotic spindle. Compared to the open space of the cell body cytoplasm, tubulin entering the cilium by diffusion will have to pass through the transition zone (TZ), a narrow opening between the cell body and the cilium (Schuss et al., 2007). The opening into the cilium corresponds to just ~0.01% of the surface of the plasma membrane and is further constricted by the TZ fibers (Tran and Lechtreck, 2015). Thus, the transition zone is a passive barrier slowing down the diffusional entry of soluble proteins into the cilium (Fig. 5). Given enough time, cell body and ciliary tubulin will equilibrate by diffusion. However, the physical barrier of the TZ could render diffusion insufficient to replace tubulin depleted from the ciliary matrix during rapid axonemal elongation in a timely manner. A transient decrease in the concentration of soluble tubulin could trigger depolymerization of the axonemal microtubules. We noted previously that the concentration of soluble tubulin inside growing cilia exceeds that of non-growing cilia even when both are present on the same cell body; the increase correlates with high frequency transport of tubulin by IFT into the growing cilium (Craft et al., 2015). Rather than having an auxiliary role in tubulin delivery, motor-based IFT could maintain the ciliary concentration of tubulin above a threshold critical for efficient axonemal assembly. IFT unloads tubulin mostly near the ciliary tip, where the tubulin concentration is particularly critical for ciliary growth and where a high tubulin concentration will be maintained longer since dilution by diffusion is most limited near the ciliary tip (Takao and Kamimura, 2017). Even with the bulk of axonemal tubulin entering cilia by diffusion, ciliary assembly could still be incapacitated in the absence of functional IFT because the intraciliary tubulin concentration will not reach the level necessary for assembly by diffusion alone. We propose that IFT regulates the intraciliary concentration of tubulin to promote axonemal assembly and cilia elongation.

**Figure 5).**
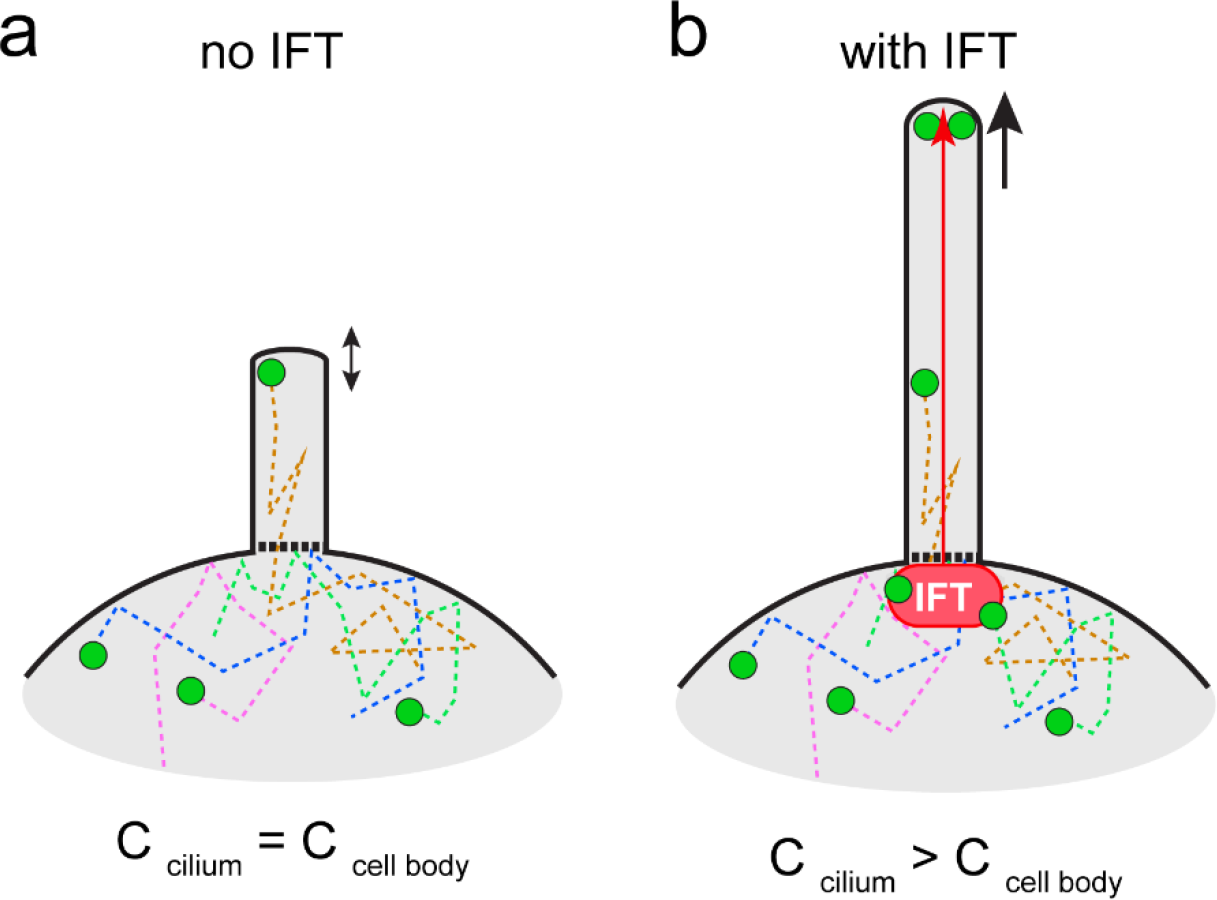
Tubulin transport by IFT enables ciliary elongation. Schematic presentation of tubulin influx into cilia without (a) and with (a) IFT. a) In the absence of IFT, tubulin (green dots) diffusing in the cell body (dashed lines) will eventually enter the cilium. Proteins entering cilia by diffusion will have to pass through the narrow openings of the transition zone at the ciliary base. This geometry will delay diffusional entry of tubulin into the cilium and axonemal elongation could decrease the concentration of soluble tubulin halting further elongation. b) IFT trains will pick-up tubulin near the flagellar base, move it through the physical barrier of the transition zone, and release it near the ciliary tip. IFT increases the tubulin concentration in the cilium, particularly near the ciliary tip, above that of the cell body promoting axonemal elongation.

## Acknowledgements

We thank Joel Rosenbaum (Yale University) for the kind gift of anti-*Chlamydomonas*-β-tubulin). This study was supported by a Uehara Memorial Foundation Research Fellowship for Research Abroad (to T.K.), the Robert W. Booth Endowment at the University of Massachusetts Medical School (to G.W.), and grants by the National Institutes of Health (R37GM030626 to G.W. and R01GM110413 to K.L.). The content is solely the responsibility of the authors and does not necessarily represent the official views of the National Institutes of Health. The authors declare that they have no conflict of interest.

## Material and Methods

### Strains and culture conditions

*C. reinhardtii* was maintained in modified M medium at room temperature, aeration with 0.5% CO_2_, and a light/dark cycle of 14:10 h. Strain CC-620 was used for most experiments (http://www.chlamycollection.org/).

### Generation of transgenic strains

The TUB2 gene encoding β-tubulin was synthesized omitting the second intron and introducing flanking BamH1 and EcoR1 sites (Genewiz). The TUA1 gene in the previously described vector pBR25-sfGFP-α-tubulin (Craft et al., 2015; Rasala et al., 2013) was replaced with the modified TUB2 using digestion with BamH1 and EcoR1 and ligation. Changes in the E-hook of β-tubulin were prepared as follows: gene segments encoding the modified C termini (E-hooks) of β-tubulin were synthesized (Genewiz), excised with EcoRI and EcoRV, and ligated into pBR25-sfGFP vector digested with the same enzymes taking advantage of a unique EcoRV site near the 3’-end of the TUB2 gene. Plasmids were restricted with KpnI and XbaI, and a fragment encompassing the functional Ble::GFP-β-tubulin cassette was gel-purified and transformed into *C. reinhardtii* by electroporation. TAP plates with 10 μg/ml zeocin were used to select transformants. Transformant colonies were transferred to liquid media in 96-well plates and screened by TIRF microscopy for expression of GFP. Expression of GFP-α-tubulin has been previously described (Craft et al., 2015). The *ift20-1* IFT20-mCherry strain was transformed with the construct encoding sfGFP-β-tubulin to obtain a double-tagged strain (Lechtreck et al., 2009).

### Isolation and fractionation of cilia

To isolate cilia for biochemical analysis we followed the protocol by (Witman, 1986). Cells were concentrated by centrifugation (1,150 × g, 3 minutes, RT) and washed with 10 mM Hepes, pH 7.4. Cells were resuspended in HMS (10 mM Hepes, 5 mM MgSO4, and 4% sucrose) and deciliated by the addition dibucaine (final concentration of 4.17 mM; Sigma-Aldrich) and vigorous pipetting. The cell bodies were removed by centrifugation and cilia were sedimented (17,000 g, 4°C, 20 min). Isolated cilia were resuspended in a microtubule-stabilizing buffer, HMEK (30 mM HEPES, 5 mM MgSO4, 0.5 mM EGTA, and 25 mM KCl) plus protease inhibitor (P9599; 1:100; Sigma-Aldrich) and demembranated by addition of NP-40 Alternative (1% final concentration; EMD Millipore) on ice for 20 min. Axonemes were pelleted from membrane/matrix fraction by centrifugation (30,000 g, 4°C, 20 min) and fractions were analyzed by SDS-PAGE and Western blotting.

### Western blotting

Ciliary proteins were separated by SDS-PAGE and transferred to PVDF membrane (Immobilon; Millipore) using standard protocols. The following primary antibodies were used: rabbit anti-Chlamydomonas-β-tubulin (1:2,000; (Silflow and Rosenbaum, 1981), mouse anti-IC2 (1:50; (King and Witman, 1990), rabbit anti-GFP (1:500; Invitrogen), and rabbit anti-NAB1 (1:5,000; Agrisera). Western blots were developed using anti-mouse or anti-rabbit secondary antibodies conjugated to horseradish peroxidase (Molecular Probes) and chemiluminescence substrate (SuperSignal West Dura; Thermo Fisher Scientific). A ChemiDoc MP imaging system was used for imaging and Image Lab (both Bio-Rad Laboratories) was used for signal quantification via densitometry.

### Ciliary regeneration

Cells were washed and resuspended in M media, deflagellated by a pH shock (pH ~4.2 for 45 s), transferred to fresh M medium, and allowed to regrow cilia under constant light with agitation (Lefebvre, 1995). To delay the onset on regeneration, cells were kept on ice until needed. To initiate regeneration, cells were diluted into ambient temperature M medium.

### In vivo microscopy

For in vivo imaging of GFP-tubulin by through-the-objective TIRF illumination, we used a Nikon Eclipse Ti-U equipped with 60× NA1.49 objective and 40-mW 488-nm and 75-mW 561nm diode lasers (Spectraphysics; (Lechtreck, 2013; Lechtreck, 2016)). The excitation lasers were cleaned up with a Nikon GFP/mCherry TIRF filter and the emission was separated using an image splitting device (Photometrics DualView2 with filter cube 11-EM). Specimens were prepared as follows. Cells (~ 10 μl) were placed inside a ring of vacuum grease onto a 24×60mm No. 1.5 cover glass and allowed to settle (~1 – 3 minutes). The observation chamber was closed by inverting a 22×22 mm No. 1.5 cover glass with ~10 μl of 5 mM Hepes, 5 mM EGTA pH 7.3 onto the larger cover glass. Images were recorded at 10–31 fps using an iXON3 (Andor) and the NIS-Elements Advanced Research software (Nikon). ImageJ (National Institutes of Health) with the LOCI plugin (University of Wisconsin, Madison WI) and multiple kymogram plugin (European Molecular Biology Laboratory) were used to generate kymograms as previously described (Lechtreck, 2013). To photobleaching the cilia, the intensity of the 488nm laser was increased to ~10% for 4 – 12 s.

### Video analysis

IFT transport events were identified manually in ImageJ. The angle tool was used to determine the velocity of the transport events. Data were transferred to Excel for further analysis. If not stated differently in the figure legend, the n values states the number of cilia analyzed. Individual frames and kymograms were save in ImageJ, Adobe Photoshop was used to adjust brightness and contrast, and figure were mounted in Illustrator.

